# Genomics reveals the origins of ancient specimens

**DOI:** 10.1101/752121

**Authors:** Qian Cong, Jinhui Shen, Jing Zhang, Wenlin Li, Lisa N. Kinch, John V. Calhoun, Andrew D. Warren, Nick V. Grishin

## Abstract

Centuries of zoological studies amassed billions of specimens in collections worldwide. Genomics of these specimens promises to rejuvenate biodiversity research. The obstacles stem from DNA degradation with specimen age. Overcoming this challenge, we set out to resolve a series of long-standing controversies involving a group of butterflies. We deduced geographical origins of several ancient specimens of uncertain provenance that are at the heart of these debates. Here, genomics tackles one of the greatest problems in zoology: countless old, poorly documented specimens that serve as irreplaceable embodiments of species concepts. The ability to figure out where they were collected will resolve many on-going disputes. More broadly, we show the utility of genomics applied to ancient museum specimens to delineate the boundaries of species and populations, and to hypothesize about genotypic determinants of phenotypic traits.

---

A study of an animal starts from its name, which is governed by strict rules^1^. A specimen termed the name-bearing type represents a species^1^. The lectotype specimen^1^ of *Homo sapiens* is Carl Linnaeus, the father of taxonomy^2^. Populations conspecific with the type carry its name. If two types are conspecific, the earlier name is used. Because most animals were named over a century ago, their types are old and lack details about their collection localities. Historically, phenotypic similarity to the type and its geographic locality determined conspecificity. For cryptic species and uncertain type localities, the phenotypic approach is problematic, leading to heated debates^3-7^. We devised a strategy to obtain genomic sequences of old type specimens and determine to which present-day populations they correspond. To solve a daunting biological problem, we applied this strategy to the skipper butterfly *Hesperia comma* and its relatives. It is *the* most important skipper, because the whole family of butterflies (Hesperiidae, the skippers) is typified by the genus *Hesperia*, and the type of that genus is *H. comma*, as designated by Carl Linnaeus himself. Several mysteries surround these butterflies. (1) Like the Linnaean type of *comma*, the type of its American counterpart, *Hesperia colorado*, collected by noted naturalist Theodore Mead^3,4^, lacks a locality label. Where was this type collected? The answer will seal the fate of names later given to populations of this species. (2) Are American *comma*-like butterflies the same species as *comma* in Europe? (3) Unusually for butterflies, *Hesperia* inhabits a wide range of elevations, from lowlands to alpine zone at over 3500 m. What are the genetic determinants of this elevational plasticity? Genomic data provide clear answers to all these questions.

## *Hesperia colorado* lectotype mapped to Lake County in Colorado

The biggest puzzle and controversy is the origin of the *H. colorado* lectotype specimen. It was collected by Mead in 1871 in Colorado, but where exactly? This question is critical because the answer determines what names apply to *Hesperia* populations. Some^5,6^ suggested that the lectotype is from the subalpine zone, while others^3,4,7^ argued that it is from the Arkansas River Basin. It was even proposed to be a hybrid of these two populations, and thus a poor choice for the name-bearing type^5,6^. Therefore, phenotypic traits without precise locality data are insufficient to attribute the lectotype to a population. Such situations are common in zoology, and offering a general solution, we ventured to place the lectotype on the map using genomics.

First, we sequenced and assembled a reference genome of *Hesperia colorado* from a specimen collected and preserved for that purpose. The genomic sequence was about 650 million base pairs, larger than most butterflies. Second, we overcame the challenge of sequencing a 150-year-old small butterfly pinned in a collection without damaging such an invaluable specimen. This resulted in ∼25% complete nuclear and entire mitochondrial genomes of the *colorado* lectotype. Third, we obtained a whole genome shotgun of 85 specimens across Colorado (Table S1). The ages of these specimens varied from 150 years (from Mead’s same expedition) to recently collected. These sequences were mapped onto the reference genome and analyzed by a combination of population genetics tools. The result was unambiguous: the lectotype of *colorado* was geographically placed in an area about 15 km in diameter in Lake County, Colorado, and belongs to the Arkansas River Basin population (Fig. 1, specimen #45).

**Fig. 1.**
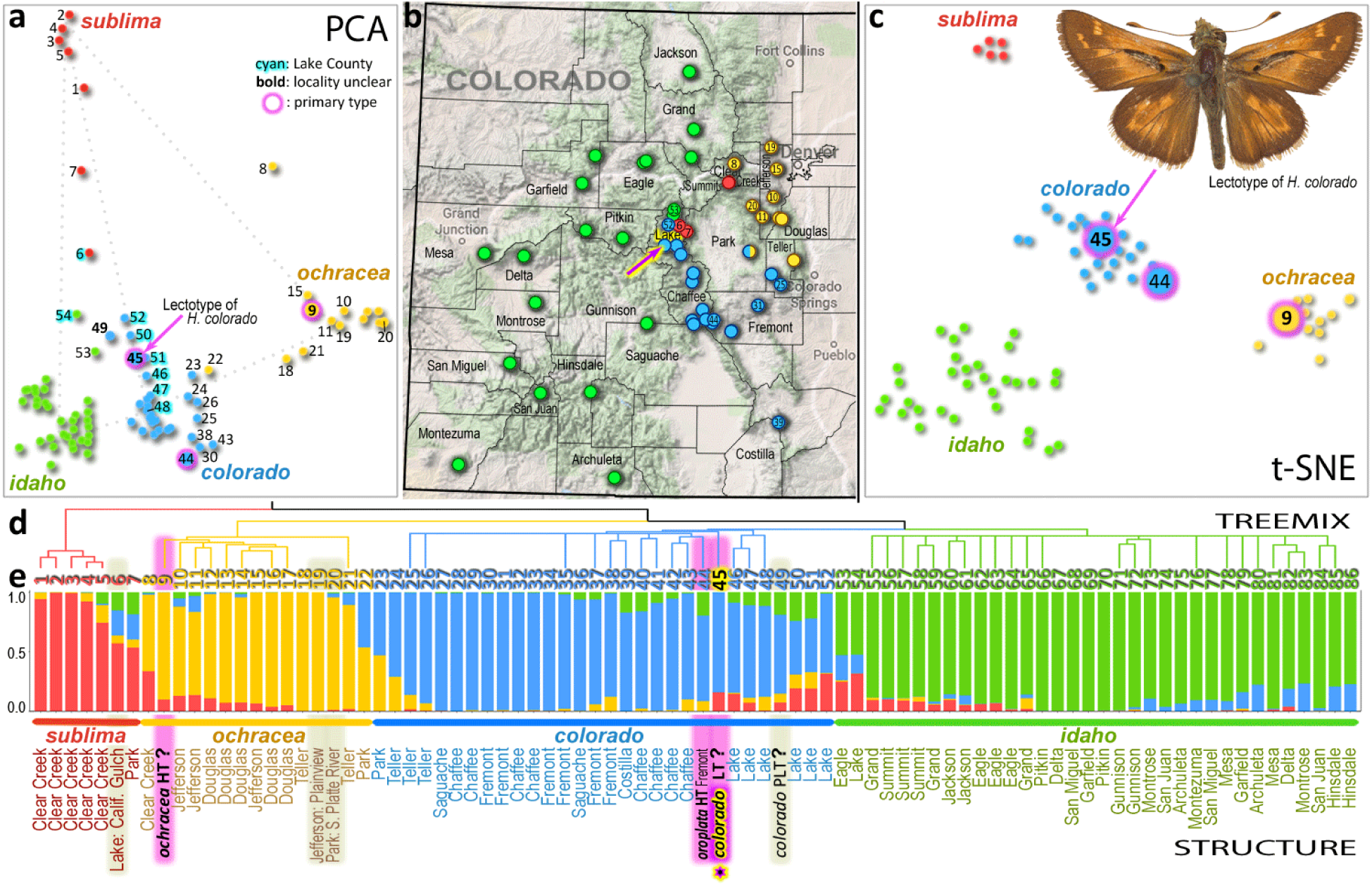
*Hesperia colorado* type specimen traced to Lake County in Colorado. Different methods consistently partition specimens into four populations: *sublima* (red), *ochracea* (orange), *colorado* (blue), and *idaho* (green). **(a)** Principal Component Analysis (PCA) of covariance between the SNPs in samples from Colorado using Eigensoft^26^. The 1st and 2nd components are shown. **(b)** Truncated map of Colorado showing specimen localities. Arrow points to the locality in Lake County we deduced for the *colorado* lectotype. **(c)** t-SNE (parameter: perplexity = 10) reduced the first 10 PCA dimensions to two, revealing populations as clusters. **(d)** TREEMIX results showing the clustering and evolutionary history of specimens. **(e)** Population structure inferred by STRUCTURE, showing the proportion of each population’s features in each specimen. Names of taxa (*colorado, oroplata, ochracea*) are given for type specimens (HT holotype, LT lectotype, PLT paralectotype), and primary types are highlighted in magenta; other ancient specimens are highlighted in gray; county names are given for other specimens (see other data in Table S1, referred to by the numbers from 1 to 86 given as leaves in the TREEMIX tree); a question mark after the name of a type indicates a questionable collection locality that we identify here using genomic comparisons.

Using PCA, t-SNE, STRUCTURE and TREEMIX analyses, we determined that *H. colorado* is represented by 4 major populations in Colorado (blue, green, red and yellow in Fig. 1). These populations correspond to subspecies. The alpine subspecies *sublima* is surrounded by three others along the three major river basins: the Platte (*ochracea*), the Arkansas (*colorado*) and, the largest, Colorado (assigned to *idaho*) (Fig. 1b). The subspecies intergrade at the boundaries of their ranges, forming hybrids (Fig. 1e). Hybridization patterns are best revealed by the PCA analysis, where the hybrids line up between the centers of populations (along dotted lines in Fig. 1a). The transition between the alpine (red) and Arkansas River Basin (blue) populations is in Lake county (cyan numbers in Fig. 1a), where three subspecies meet and hybridize. The lectotype of *colorado* (#45 Fig. 1a) is surrounded exclusively by the specimens from Lake Co., implying where it was collected. Moreover, the mitogenome of the lectotype was a 100% match to a single specimen (Fig. 2c), and that specimen was collected near Twin Lakes in Lake Co. Another of Mead’s specimens (#49, a paralectotype, Fig. 1a), collected during the same 1871 expedition, maps to the same area. STRUCTURE (Fig. 1e) reveals their genomic composition. Lake County specimens are characterized by having some *sublima* (red) and *idaho* (green) components, making them attributable to the narrow geographic zone of transition between these populations. Nevertheless, as t-SNE shows (Fig. 1c), the lectotype clusters within the Arkansas River Basin population (called *oroplata*; #44 is its holotype), and STRUCTURE reveals that only a minor fraction of its genome is of hybrid origin. Thus the name for this population is *colorado*, superseding *oroplata*, a younger name^8^.

**Fig. 2.**
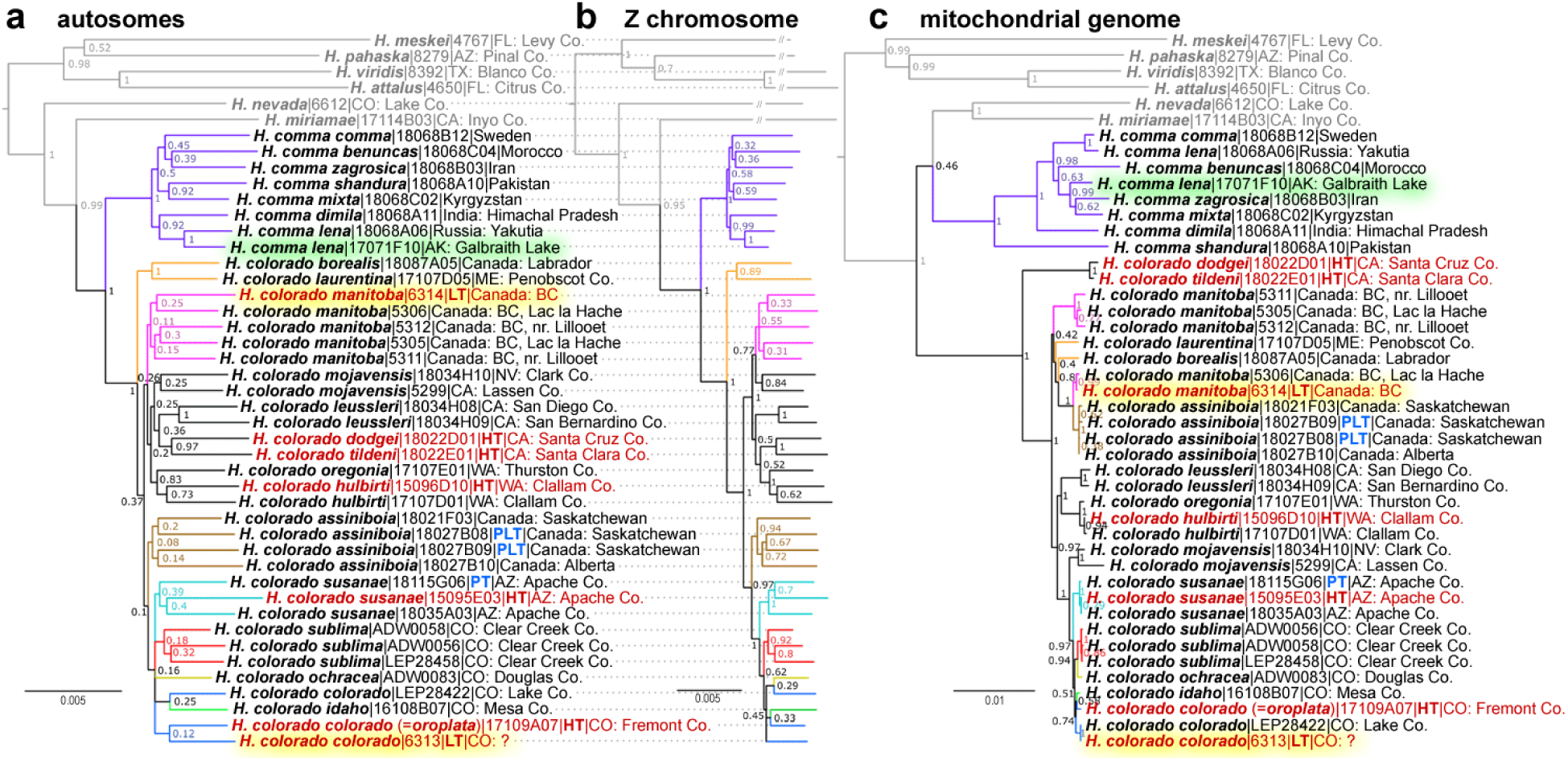
Phylogenetic analysis suggests that *Hesperia colorado* is a species distinct from *Hesperia comma.* The trees are based on concatenated genomic alignments from **(a)** autosomes, **(b)** Z chromosome, and **(c)** mitogenome. Name, voucher number and general locality are given for each specimen, see Table S1 for additional data. *Hesperia* species outside the *comma* group are shown in gray. The trees are rooted with *Pompeius verna* (NVG-18014H01, not shown). Names of primary type specimens are shown in red. The two types with questionable localities are highlighted in yellow and an American *H. comma* specimen from Alaska is highlighted in green.

As a control of our methods applied to ancient specimens, we used three others (two collected by Mead) with less questionable localities^3,4^. These specimens (#6, 19 & 20, Fig. 1) mapped accordingly with their presumed localities. Specimen #6, from near Leadville, is indeed placed with the Lake Co. specimens, but closer to *sublima* as suggested by its genetic makeup. The other two specimens were attributed to *ochracea* in agreement with their associated data. Placing these controls builds the confidence in inferring localities of specimens #45 & 49 by genomics. Moreover, historical evidence suggests that the lectotype was indeed collected in Lake County, as the specimen bears a label “7-13” (July 13). According to Mead’s journal, he was at Twin Lakes on that date. We think that our success in deducing the localities is due to *Hesperia* being non-migratory, local butterflies that form well-diverged populations.

Another type specimen with a questionable collection locality, *H. colorado manitoba*, trees next to a specimen from Lac la Hache in British Columbia (Fig. 2a-c), in accordance with its label data, thereby validating *manitoba*’s type locality^9^. Topology of the nuclear genomic trees follows the geographic distribution of these populations: those that are close on the map tend to cluster in the trees. For example, all specimens from Colorado, including the *colorado* lectotype, form a clade, which also includes the nearby populations from eastern Arizona. However, the mitogenome tree (Fig. 2c) is incongruent with the nuclear genome trees, revealing a history of mitochondria different from that of nuclear genomes, a phenomenon commonly observed in closely related populations and species^10^. Nevertheless, all three trees support the distinction between Old World *comma* (also found in Alaska) and the North American species *H. colorado*, and place both *manitoba* and *colorado* type specimens in agreement with their collection localities^3,9^, contrary to speculations by Scott et al^5,6^.

## *Hesperia colorado* is a species distinct from *Hesperia comma*

Many American species have similar-looking counterparts in Europe and Asia, posing a question about whether we should regard them as the same species. Phylogenetic trees of three different genomic regions consistently reveal a deep split between American *Hesperia colorado* and the Old World *Hesperia comma* (Fig. 2), strongly suggesting that they are not conspecific. The genome-wide Fixation index (*Fst*)^11^ between *comma* and *colorado* is about 0.5, corresponding to a divergence between species. Despite its broad distribution, populations of *comma* from Europe, Africa and Asia are much closer to each other than any of them to Nearctic *colorado*. A surprise was a specimen from northern Alaska, which is true *comma*, most similar to specimens from northeastern Russia across the Bering Strait (Fig. 2ab). Therefore, *comma* is indeed present in America, but all other *comma*-like specimens we sequenced from the US and Canada were *colorado*. Functional analysis (Table S2) of proteins that differ the most between the two species reveals the prevalence of nuclear-encoded mitochondrial respiratory chain components, which is consistent with the profound divergence (about 3%) in the mitogenomes. Furthermore, the two species show divergence in circadian rhythm, DNA, and histone methylation, which may have a direct effect on the reproductive barrier between *comma* and *colorado*.

## Adaptations to high elevation in *Hesperia colorado sublima*

Unlike other *Hesperia colorado* populations, *sublima* is confined to high elevations. Comparing *sublima* with others, we identified 162 unique amino acid replacements enriched in 80 proteins and positively selected in 7 (Table S3). These proteins function in metabolism, response to light, and development of the respiratory system and muscles (Fig. 3ab). This diversity, more than in human high-elevation populations dated up to 45,000 ya^12^, is likely due to the longer time of adaptation (∼100,000 ya) and one to two year generation span in *Hesperia*. Human adaptations are largely restricted to variations in hypoxia-inducible factors, which upregulate glycogen and ATP synthesis under hypoxic conditions. Similarly, in *sublima*, variations map to regulators of insulin pathway and glycogen synthase (GS), in addition to metabolic enzymes, electron carriers and heme-binding proteins. These variations may reprogram *sublima* metabolism to ensure sufficient energy production with reduced oxygen. One GS variation is in the helix interacting with glycogenin (Fig. 3c). The interaction between the GS and glycogenin is crucial^13^, and the *sublima*-specific variation affecting this interaction may lead to more efficient glycogen production.

**Fig. 3.**
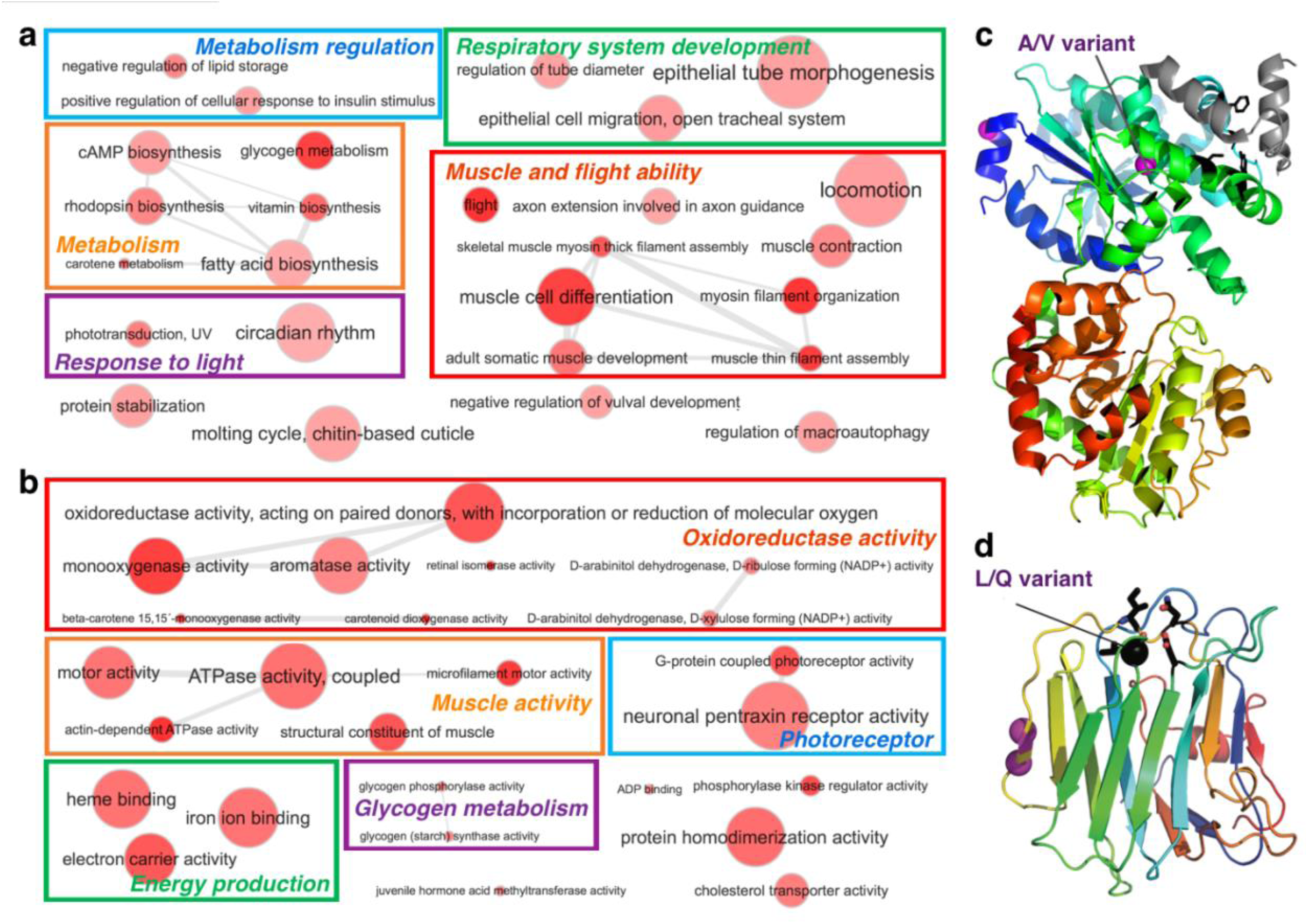
Molecular adaptation to high elevation in *Hesperia colorado sublima*. **(a-b)** Functions of proteins related to high-elevation adaptation revealed by Gene Ontology (GO) terms in the categories of **(a)** biological process and **(b)** molecular function. The size of a circle correlates with the number of proteins in the genome associated with that GO term and its color indicates the significance (darker color corresponds to lower P-value) of a GO term enrichment among proteins showing high-elevation adaptation. **(c)** 3D structure model of glycogen synthase (template PDB: 4QLB. A variant in *sublima* (from A to V) is near the functional sites (black sticks) mediating its interaction with glycogenin. **(d)** 3D structure model (template PDB:3POY) of motor neuron guidance factor *trol* with a variant (black sphere, L in *sublima*, and Q in low-elevation populations) near the Ca2+ binding site. Other variants are shown as magenta spheres.

Insects lack the dedicated oxygen-carrying blood cells of vertebrates, and a network of tracheal tubes directly delivers oxygen throughout the body. Similar to humans growing more capillaries in muscle with exercise at high elevation^14^, *sublima*-specific variations in factors governing the development of this network may promote elaboration of tracheal tubes. Other adaptations in *sublima* lack apparent human analogies. We observe variations in UV photoreceptors and circadian regulators, changes likely related to stronger UV radiation and colder climate at higher altitudes. Most notably, 4 out of 7 proteins positively selected in *sublima* are related to flight ability, including the motor neuron axon guidance factor *trol* with a variation near its calcium binding site: in *sublima*, hydrophobic Leucine replaces hydrophilic Glutamine of lower elevation populations (Fig. 3d). Strong fliers, including swallowtails and monarch butterflies, all have hydrophobic residues in this position, while in weak fliers, such as the cabbage white, this residue is hydrophilic. These changes should make *sublima* a better flier adapted to stronger wind on mountaintops.

## Genetic basis for paler appearance of *Hesperia colorado ochracea*

Out of all populations in Colorado, *ochracea* is perhaps most recognizable due to its paler wings and poorly defined white spots. Surprisingly, we found that *ochracea*-specific polymorphisms are largely confined to a short 200kb region on the Z chromosome (Fig. 4). This unusually high density of SNPs suggests introgression from some population outside of Colorado rather than gradual evolution by point mutations. ABBA-BABA tests^15^ with other populations reveal that this region was introgressed (P-value < 6e-66) from *Hesperia colorado assiniboia*, a northern butterfly of the Central Plains. Because the wing patterns of *ochracea* resemble *assiniboia* (Fig. 4a-f), it is likely that this introgressed region is inducing *assiniboia-*like wing patterns in *ochracea.* Proteins encoded by this region include Shank3, a regulator of Wnt signaling pathway modulating internalization of the Wnt receptor Fz2^16^. The Wnt pathway has been implicated in wing patterning^17-19^, and Wnt receptor Fz2 is expressed in developing wings^20^, suggesting a role of Shank3 in wing pattern formation.

**Fig. 4.**
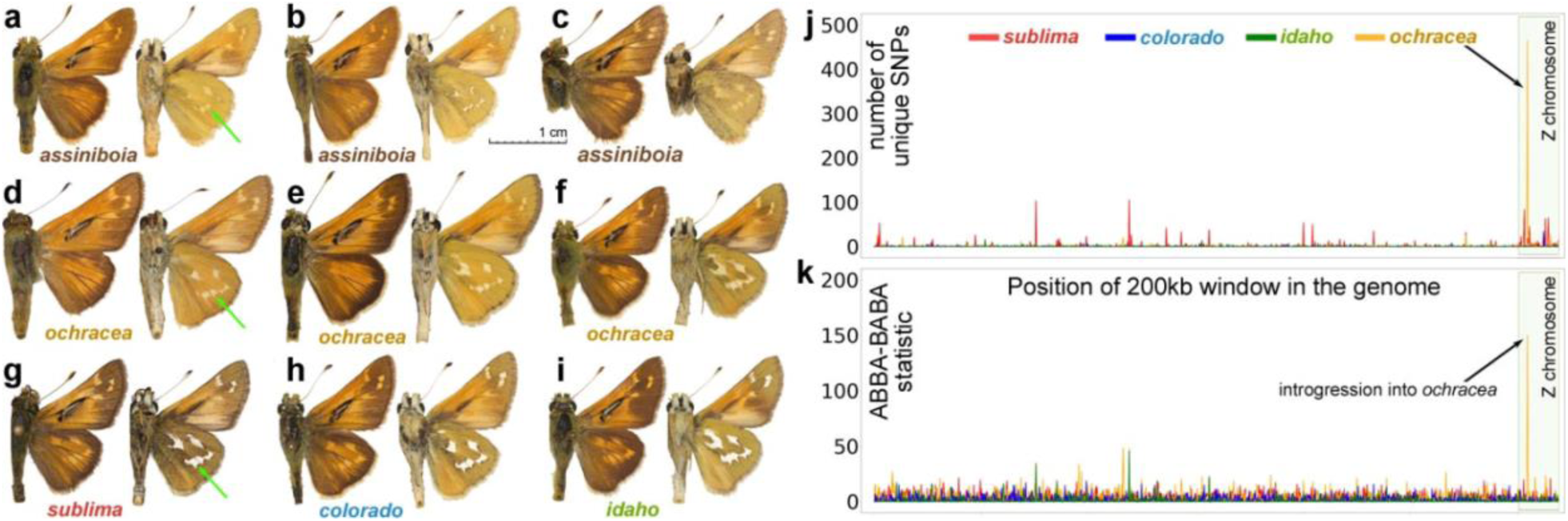
Similarity in wing pattern between *Hesperia colorado ochracea* and *H. c. assiniboia* likely caused by the introgression of a 200kb Z-linked genomic region. Wing pattern variation in **(a-c)** *assiniboia* and **(d-f)** *ochracea* specimens; **(g-i)** typical specimens of other *Hesperia* populations in Colorado. Green arrows denote pale spots on the hindwing differing between populations. Voucher numbers for (a-i) are NVG-18027B08, NVG-18027B09, NVG-18027B10, NVG-15111B01, NVG-5533, NVG-16108A05, NVG-5532, NVG-16108C02, and NVG-16108B04, respectively; see Table S1 for additional data. **(j)** Number of unique SNPs in different populations in 200kb windows throughout the genome. **(k)** Introgression from *assiniboia* to 4 *Hesperia* populations in Colorado identified using the ABBA-BABA test. The y-axis shows the difference between the number of positions with the pattern ABBA and the number of positions with the pattern BABA in a 200kb genomic window (negatives omitted). The x-axis on both plots is the position of the window in the genome (concatenated scaffolds). Z chromosome scaffolds are placed last (highlighted pale olive). Counts are colored by taxon: *sublima* (red), *colorado* (blue), *idaho* (green), and *ochracea* (orange).

## Materials and methods

Specimens used in this project were either collected in the field (and stored in RNAlater or EtOH), or borrowed from collections listed in the Acknowledgements. The collection dates of specimens ranged from 1871 to 2016. See Table S1 for complete specimen data. A piece of thoracic tissue from fresh specimens, and either the abdomen or a leg from pinned museum specimens were used for DNA extraction and genomic library preparation according to our protocols developed previously^21-24^. We used mate-pair libraries to assemble a reference genome of *Hesperia colorado* from a single wild-collected specimen, also accompanied by RNAseq for gene annotation. Paired-end libraries were constructed for all other specimens, sequenced for 150 bp from both ends targeting 5 to 10X coverage and mapped to the reference genome. Z chromosome scaffolds were found as those containing Z chromosome proteins in *Heliconius*^25^. The following computational tools were used: PCA^26^, t-SNE^27^, TREEMIX^28^, STRUCTURE^29^, and IQ-TREE^30^. Details of adapting these methods to work with data from ancient specimens are described in the Supplement.

## Acknowledgments

We are indebted to Naomi Pierce, Philip Perkins and Rachel Hawkins (Museum of Comparative Zoology, Harvard University) for the loan of historically significant *Hesperia* specimens for DNA analysis making this work possible. We are grateful to David Grimaldi and Courtney Richenbacher (American Museum of Natural History, New York, NY), Jonathan Pelham (Burke Museum of Natural History and Culture, Seattle, WA), John Rawlins (Carnegie Museum of Natural History, Pittsburgh, PA), Paul Opler and Boris Kondratieff (Colorado State University Collection, Fort Collins, CO), Weiping Xie (Los Angeles County Museum of Natural History, Los Angeles, CA), Rodolphe Rougerie (Muséum National d’Histoire Naturelle, Paris, France), Edward Riley, Karen Wright, and John Oswald (Texas A & M University, College Station, TX), Robert Robbins, John Burns, and Brian Harris (National Museum of Natural History, Smithsonian Institution, Washington, DC) for granting access to the collections under their care and for stimulating discussions; Ernst Brockmann, Steve Kohler, and Mark Walker for sampling and providing photos of specimens in their collections; Texas Parks and Wildlife Department (Natural Resources Program Director David H. Riskind) for the permit #08-02Rev that makes research based on material collected in Texas State Parks possible. The study has been supported by grants from the National Institutes of Health GM127390 and the Welch Foundation I-1505 to NVG.

